# Behavioral evidence for memory replay of video episodes in macaque monkeys

**DOI:** 10.1101/2020.01.10.902130

**Authors:** Shuzhen Zuo, Lei Wang, Junghan Shin, Yudian Cai, Sang Wan Lee, Kofi Appiah, Yong-di Zhou, Sze Chai Kwok

## Abstract

Humans recall the past by replaying fragments of events temporally. Here, we demonstrate a similar effect in macaques. We trained six rhesus monkeys with a temporal-order judgement (TOJ) task and collected 5000 TOJ trials. In each trial, they watched a naturalistic video of about 10 s comprising two across-context clips, and after a 2-s delay, performed TOJ between two frames from the video. The monkeys apply a non-linear forward, time-compressed replay mechanism during the temporal-order judgement. In contrast with humans, such compression of replay is however not sophisticated enough to allow them to skip over irrelevant information by compressing the encoded video globally. We also reveal that the monkeys detect event contextual boundaries and such detection facilitates recall by an increased rate of information accumulation. Demonstration of a time-compressed, forward replay like pattern in the macaque monkeys provides insights into the evolution of episodic memory in our lineage.

**Impact Statement:** Macaque monkeys temporally compress past experiences and use a forward-replay mechanism during judgment of temporal-order between episodes.

## INTRODUCTION

Accumulating evidence indicate that non-human primates possess the ability to remember temporal relationships among events (*1*–*3*). The apes can remember the movies based on the temporal order of scenes (*4*) and keep track of past time of episodes (*5*), whereas macaque monkeys possess serial (*6*) and ordinal positions expertise for multi-item lists (*7*) and are able to categorize sequences of fractal images by their ordinal number (*8*). However, keeping track of and remembering the positional coding and forming associative chaining (*9*, *10*) of lists of arbitrary items might diverge from how a semantically linked, temporally relational representation of real-life events is maintained and utilized.

In the human literature, it has been shown that episodes can be replayed sequentially based on learned structures (*11*), sensory information (*12*), and pictorial content (*13*). These findings suggest the possibility that monkeys can rely on a similar mechanism in recalling events that are linked temporally during temporal order judgement (TOJ). However, the extent to which mechanisms of temporal order judgement overlap across humans and monkeys remains undefined. One hypothesis is that the macaques can similarly rely on a scanning model for information retrieval – akin to serial replay of episodes – to perform temporal order judgements (*2*, *3*). By this account, following encoding streams of events, the animal performs retrieval through replaying the stream of information in a forward direction. In this way, retrieval time (RT) would be positively correlated to the temporal distance between the beginning of the stream and the target location. This account aligns with recent findings in rodents (*14*) and in humans of memory replay during cued-recall tasks across fragments of video episodes, showing characteristics of replay proceeding in a forward manner and being temporally compressed (*12*). Moreover, a more sophisticated feature of this mechanism is that memory replay is a fluidic process in which subjects are able to skip flexibly across sub-events (*12*). By this account, subjects can omit non-informative parts of episodes and replay a shorter episode (shorter than physical perception) in memory, which contains less information. This interpretation is supported by other works on mental simulation of paths (*15*) and video episodes (*12*). This latter account constitutes a global compression of parts of episodes (that allows skipping across sub-events) and is regarded as substantially superior to a strict forward replay mechanism.

In order to simulate dynamic flow of information as in real-life scenarios, we used naturalistic videos as experimental material to study the mechanism of memory retrieval of event order in the monkeys, which are more realistic than arbitrary items or images that were used in previous studies (*1*, *16*). We used a temporal order judgement paradigm to examine whether and to what extent the pattern underlying memory retrieval conforms to a time-compressed, forward replay mechanism. In each trial, monkeys watched a naturalistic video, which is composed of two clips, and following a 2-s retention delay, made a temporal order judgement of choosing the frame that was shown earlier in the video between two frames extracted from that video (Figure 1A). The two frames were either extracted from the same clip or from two different clips of the video. Given that analyses on response latency can provide insights into the extent to which the monkeys’ behavior might conform to the two putative replay models outlined above, we looked into the RT data. By applying representational similarity analyses (RSA), the Linear Approach to Threshold with Ergodic Rate (LATER) model and generalized linear models on the RT data, we examined the presence of replay-like behavioral patterns in the monkeys. Specifically, if monkeys recall the frames by their ordinal positions, this would imply a linear increase in their retrieval times. In contrast, if the memory search entails a complex processing of the content determined by their semantically linked, temporally relational linkage within the cinematic footage, we should observe evidence for some non-linear pattern.

**Figure 1.**
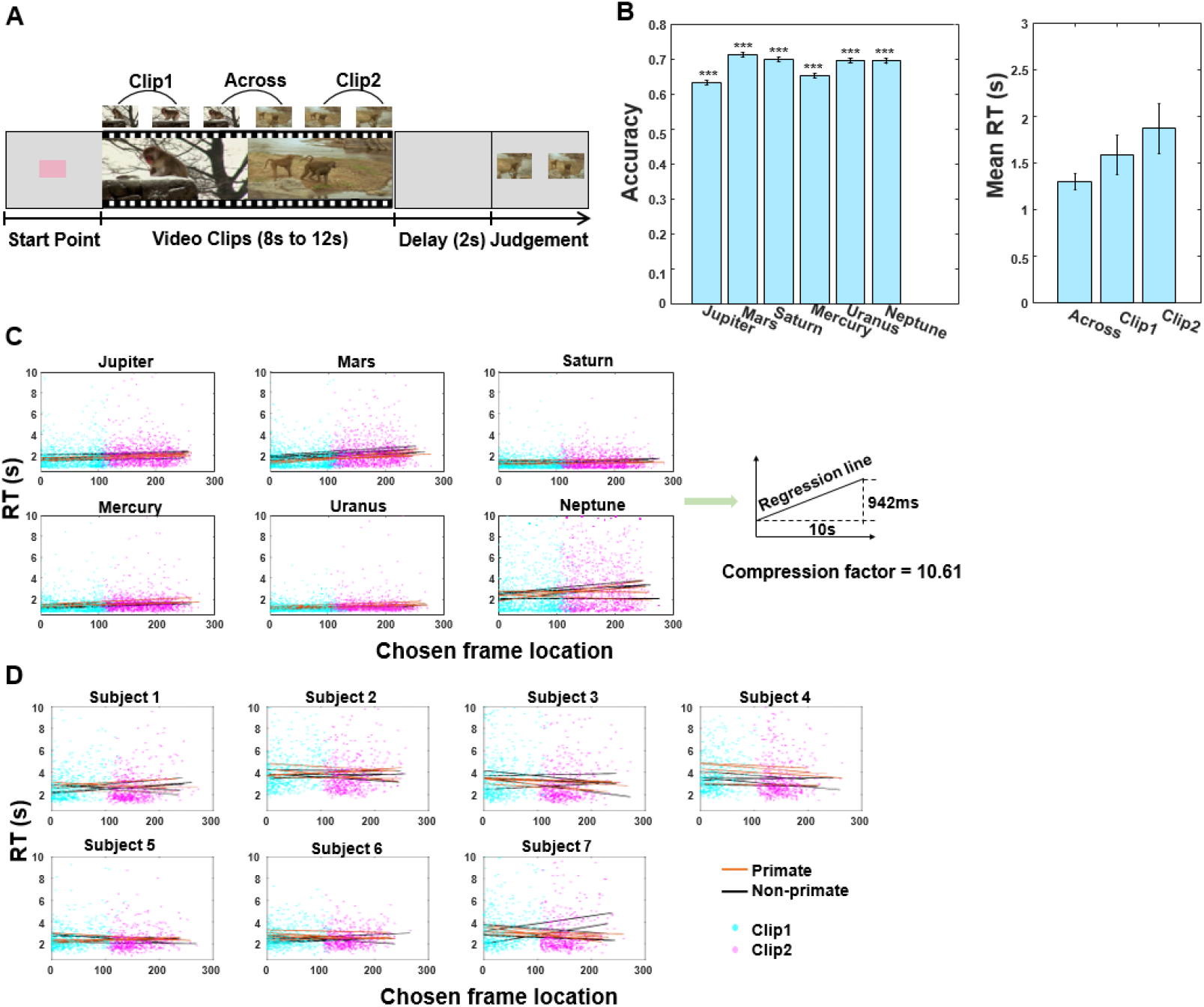
TOJ task schema and RT results. **(A)** On each trial, the monkey watched a video (8-12s, comprising two 4-6s video clips), and following a 2-s retention delay, made temporal order judgement between two probe frames extracted from the video. The monkeys were required to choose the frame that was presented earlier in the video for water reward. **(B)** Task performance of six monkeys. Proportion correct for the six monkeys (left); mean reaction times for three trial types (right). Error bars are standard error of the mean (SEM). *** denotes P < 0.001. **(C)** Linear regression analysis of reaction time (RT) as a function of chosen frame location, see also Table 1. **(D)** Linear regression analysis of RT as a function of chosen frame location for each human participant, see also Table S3. In (C) and (D), black lines and orange lines refer to lists of non-primate and primate videos respectively (with five repetitions collapsed for monkeys and two repetitions collapsed for human participants). All responses in within-context are shown, with cyan and magenta dots denoting whether the chosen probe frames were extracted from either Clip 1 or Clip 2 respectively.

Our results showed that macaque monkeys adopt a time-compressed, replay-like pattern to search within representation of continuous information. In addition, by comparing between human and macaque data, we adjudicated between the two aforementioned aspects of the replay models. The results showed a discrepancy between the two species that the monkeys do not compress the cinematic events globally as effectively as in humans. Finally, we revealed that the monkeys can make use of context changes to facilitate the memory retrieval by an increased rate of information accumulation in a drift diffusion model framework.

**Table 1.** One sample t-tests results of the slopes of RT as a function of chosen frame location for each monkey. The three panels correspond to analyses performed using all trials (top), only correct trials (middle), and only incorrect trials (bottom). Same slope patterns were observed irrespective of correctness.

| Monkeys | Mean (SEM) | t-statistics | d.f. | p-value | Cohen's d | 95% confidence interval |  |
| --- | --- | --- | --- | --- | --- | --- | --- |
|  |  |  |  |  |  | Lower | Upper |
| Slope of reaction times/chosen frame location tested against zero (all trials) |  |  |  |  |  |  |  |
| Jupiter | 0.0015 (0.0017) | 6.06 | 49 | <0.001 | 0.86 | 0.0010 | 0.0020 |
| Mars | 0.0031 (0.0027) | 7.93 | 49 | <0.001 | 1.12 | 0.0023 | 0.0038 |
| Saturn | 0.0007 (0.0016) | 3.29 | 49 | 0.002 | 0.47 | 0.0003 | 0.0012 |
| Mercury | 0.0017 (0.0018) | 6.77 | 49 | <0.001 | 0.96 | 0.0012 | 0.0022 |
| Uranus | 0.0011 (0.0014) | 5.12 | 49 | <0.001 | 0.72 | 0.0006 | 0.0015 |
| Neptune | 0.0024 (0.0049) | 3.34 | 46 | 0.002 | 0.49 | 0.0009 | 0.0038 |
| Slope of reaction times/chosen frame location tested against zero (correct trials) |  |  |  |  |  |  |  |
| Jupiter | 0.0023 (0.0042) | 3.85 | 49 | <0.001 | 0.55 | 0.0011 | 0.0035 |
| Mars | 0.0029 (0.0034) | 5.98 | 49 | <0.001 | 0.85 | 0.0019 | 0.0038 |
| Saturn | 0.0009 (0.0021) | 3.03 | 49 | 0.004 | 0.43 | 0.0003 | 0.0015 |
| Mercury | 0.0018 (0.0026) | 4.79 | 49 | <0.001 | 0.68 | 0.0010 | 0.0025 |
| Uranus | 0.0012 (0.0021) | 4.01 | 49 | <0.001 | 0.57 | 0.0006 | 0.0018 |
| Neptune | 0.0030 (0.0070) | 2.94 | 46 | 0.005 | 0.43 | 0.0009 | 0.0051 |
| Slope of reaction times/chosen frame location tested against zero (Incorrect trials) |  |  |  |  |  |  |  |
| Jupiter | 0.0016 (0.0032) | 3.49 | 49 | 0.001 | 0.49 | 0.0007 | 0.0025 |
| Mars | 0.0044 (0.0039) | 8.00 | 49 | <0.001 | 1.13 | 0.0033 | 0.0055 |
| Saturn | 0.0011 (0.0025) | 3.20 | 49 | 0.002 | 0.45 | 0.0004 | 0.0018 |
| Mercury | 0.0022 (0.0026) | 5.96 | 49 | <0.001 | 0.84 | 0.0014 | 0.0029 |
| Uranus | 0.0012 (0.0019) | 4.52 | 49 | <0.001 | 0.64 | 0.0007 | 0.0017 |
| Neptune | 0.0034 (0.0075) | 3.15 | 46 | 0.003 | 0.46 | 0.0012 | 0.0056 |

## RESULTS

### Human-like forward replay in macaques

All six monkeys learned to perform the temporal order judgement task with dynamic cinematic videos as encoded content (Figure 1B left and **Movie S1**). Note that there are two main kinds of TOJ trials: “within-context” and “across-context” trials (Figure 1A). Here, we will be first concerned with the response times (RT) data from “within-context” trials to examine TOJ mechanisms, while RT data from “across-context” trials will be used to test for effects by context changes (event boundary) in subsequent subsections.

We first addressed our main hypothesis by examining changes in RT using only within-context trials. We regressed RT as a function of temporal similarity (TS) for within-context trials and found a significant negative relationship between TS and RT, all *P* < 0.001 (one monkey with *P* = 0.02, Table S2 upper panel and Figure S1). The results were replicated when considering only correct trials or only incorrect trials separately (Table S2 middle/bottom panels). This suggests that the monkeys are systemically faster in identifying frames that are located earlier in the video. One possibility is that the monkeys perform TOJ relying on some form of memory replay by recalling the events coded at specific ordinal positions in a serial manner. To test this hypothesis, in each trial we explicitly looked at the relationship between RT and the location of the frame within the video which the monkeys have chosen (“chosen frame location”, as indexed by the ordinal frame numbers in the video). By correlating RT as a function of chosen frame location, we indeed found a significantly positive relationship between RT and chosen frame location in all monkeys, all *P* < 0.001 (two monkeys with *P* = 0.002, Figure 1C and Table 1 top panel). The same pattern was replicated when only correct trials (Table 1 middle panel) or incorrect trials (Table 1 bottom panel) were included for analysis, suggesting that the putative retrieval RT patterns are not affected by the memory outcome. This pattern of result is also replicated using logarithmically transformed RT data, all *P* < 0.001. After taking individual variability into consideration, we obtained the same result with reciprocal latency as a function of chosen frame location for the average of all monkeys (Figure 2A). While the result suggests that the monkeys’ judgments are faster to respond to probe frames that are located at the earlier parts of the videos, it also indicates that the search does not follow a linear function. This non-linear relationship is important because it rules out alternative explanations that the main effects are simply resultant from positional effects. Rather, the non-linear change in slope indicates there are other factors in play that the monkeys might group (or parse) the content according to the relational storyline structure, and do not merely recall the frames/items as of their ordinal positions in a fixed linear manner.

**Figure 2.**
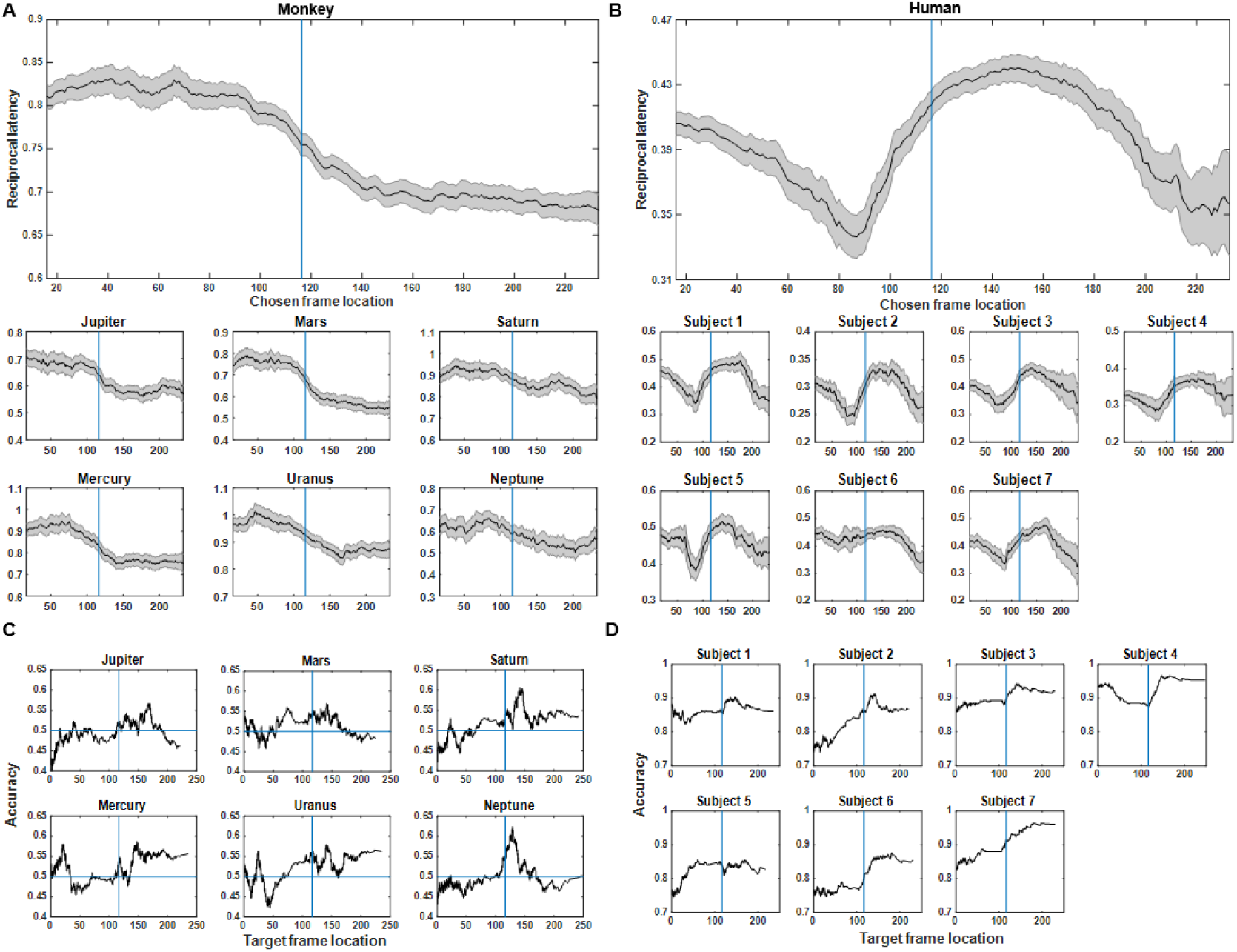
Moving average analysis based on reciprocal latency and accuracy for monkeys (left panels) and human participants (right panels). **(A)** Reciprocal latency for monkeys as a function of chosen frame location for the average of all animals (upper panel) in within-context condition, with results of six individual monkey shown (lower panel). The relationship between chosen frame location and RT follows a non-linear pattern. **(B)** Reciprocal latency for human participants as a function of chosen frame location for the average of all human subjects (upper panel) in within-context condition, with results of individual subject shown (lower panel). In (A) and (B), shaded region denotes confidence intervals; the blue line denotes the mean boundary location between Clip 1 and Clip 2 across all trials (116^th^ frame). **(C)** Individual monkey’s proportion correct as a function of target frame location in within-context condition. Horizontal blue lines denote chance level accuracy. **(D)** Individual human subject’s proportion correct as a function of target frame location in within-context condition.

To address the direction and speed of memory reply, reaction times at retrieval were compared between TOJ frames that are within Clip 1 versus Clip 2. During TOJ retrieval, frames that were presented in Clip 1 (mean reaction time = 1.59 s) were retrieved significantly faster than the frames that were experienced in Clip 2 (mean reaction time = 1.88 s) (one-tailed *t*5 = −4.533; *P =* 0.003; Cohen’s d = −0.54; 95% CI: [-inf, −0.158]; log(RT): one-tailed *t*5 = −6.473; *P* = 6.558×e^−4^; Cohen’s d = −0.69; 95% CI: [-inf, −0.114]). These findings again confirm that the replay of the video takes place in a forward direction.

Moreover, since the latency required to responding to the chosen frames is much smaller than the duration of the videos themselves, the replay of the video must have been conducted at a compressed speed (i.e., memory replay was faster relative to perception during video-watching). The difference in reaction time between the very first frame and the last frame was averaged at 942 ms (range: 468 – 1859 ms). This is equivalent to 94.2 ms to scan through per second of the video and corresponds to a compression factor of 10.61 during replay in these monkeys (compression factor for each monkey: Jupiter = 13.59, Mars = 7.39, Saturn = 14.80, Mercury = 21.37, Uranus = 17.78, Neptune = 5.38, see Figure 1C). This is comparable to a compression factor of 13.7 observed in humans, corroborating the notion of forward replay and those findings in humans (*12*).

We also ran the same sets of analyses on human participant data for comparison between the two species. Showing a completely opposite pattern, regression analyses on human subjects showed a positive relationship between temporal similarity (TS) and RT for all participants, all *P* < 0.01 (Table S3 upper panel). These results imply that the more similar the two frames to be judged, the longer time needed for retrieve temporal order information. There was also no observable RT/ chosen frame location slope in the human data (if anything, it shows an opposite trend; see Figure 1D & Table S3 bottom panel). In sum, we found that our monkeys might have performed TOJ of video episodes using a forward search of ordered elements in the mnemonic representation at the time of memory test with a non-linear compression function. By contrasting with Figure 2A, it is notable that when reciprocal latency as a function of chosen frame location is analyzed in the humans, as shown in Figure 2B, a very different pattern emerges, suggesting some form of mechanistic discrepancy between the species. We will examine these aspects in detail in the next section.

We have also run a sliding-window average analysis to illustrate how accuracy varies as a function of the target frames location (Figure 2C for monkeys; Figure 2D for humans). It is interesting that in both species there is a mild trend that their proportion correct show a little blip right after the beginning of Clip 2. This characteristic might be related to their ability in detecting the boundaries (cf. results and discussion related to Figure 6). However, given that we are primarily concerned with using response times as quantitative measures to examine the TOJ mechanisms, we do not speculate further into this aspect.

### Discrepancy with humans: Compression of replay is local but not global

It has been shown in the humans that memory replay is not a straightforward recapitulation of the original experience. Subjects can skip through their memories on a faster time scale across segments of a video episode than within-segment by skipping flexibly over salient elements such as video boundaries within episodes (*12*). We propose two possible models with respect to whether the compression is global or not over the whole video. If there is a global compression of the video during replay, the speed to initiate replay for Clip 2 would be sooner than the endpoint of replay for Clip 1 due to the animal being able to skip over the entire Clip 1 to the beginning of Clip 2 (Global-compression model, Figure 3A right panel). However, if the monkeys are not equipped with the ability of skipping video segments during the replay process, we would expect a linear increase of retrieval time with chosen frame location irrespective of the boundary (Strict forward model, Figure 3A left). We tested statistically whether the speed to initiate replay for Clip 2 slowed down compared to that for Clip 1. We divided each video into eight equal segments and derived cross-correlations computed on pairs of averaged condition-wise RTs based on chosen frame locations using a representational similarity analysis. The RT for TOJ between each segment of the video increases linearly according to their position in encoding (Figure 3B left). We tested these against a hypothetical “Strict forward” model and found significant correlation with the Strict forward model (*r* = 0.66, *P* = 0.009), but not with the Global compression model (*r* = −0.16, *P* = 0.802) (Figure 3B right). These statistics also remain significant for the Strict forward model when we divided the video into either 10 (*P* = 0.030) or 14 equal segments (*P* = 0.020). The same patterns are also obtained when considering correct trials (Strict forward model: *r* = 0.37, *P* = 0.040; Global compression model: *r* = −0.02, *P* = 0.545) or incorrect trials (Strict forward model: *r* = 0.45, *P* = 0.010; Global compression model: *r* = −0.06, *P* = 0.545) separately. Contrarily, these correlational patterns with the Strict forward model are not observed in the human subjects (*r* = −0.11, *P* = 0.703), but rather we observe a trend favoring the global compression model instead (*r* = 0.39, *P* = 0.069) (Figure 3B right).

**Figure 3.**
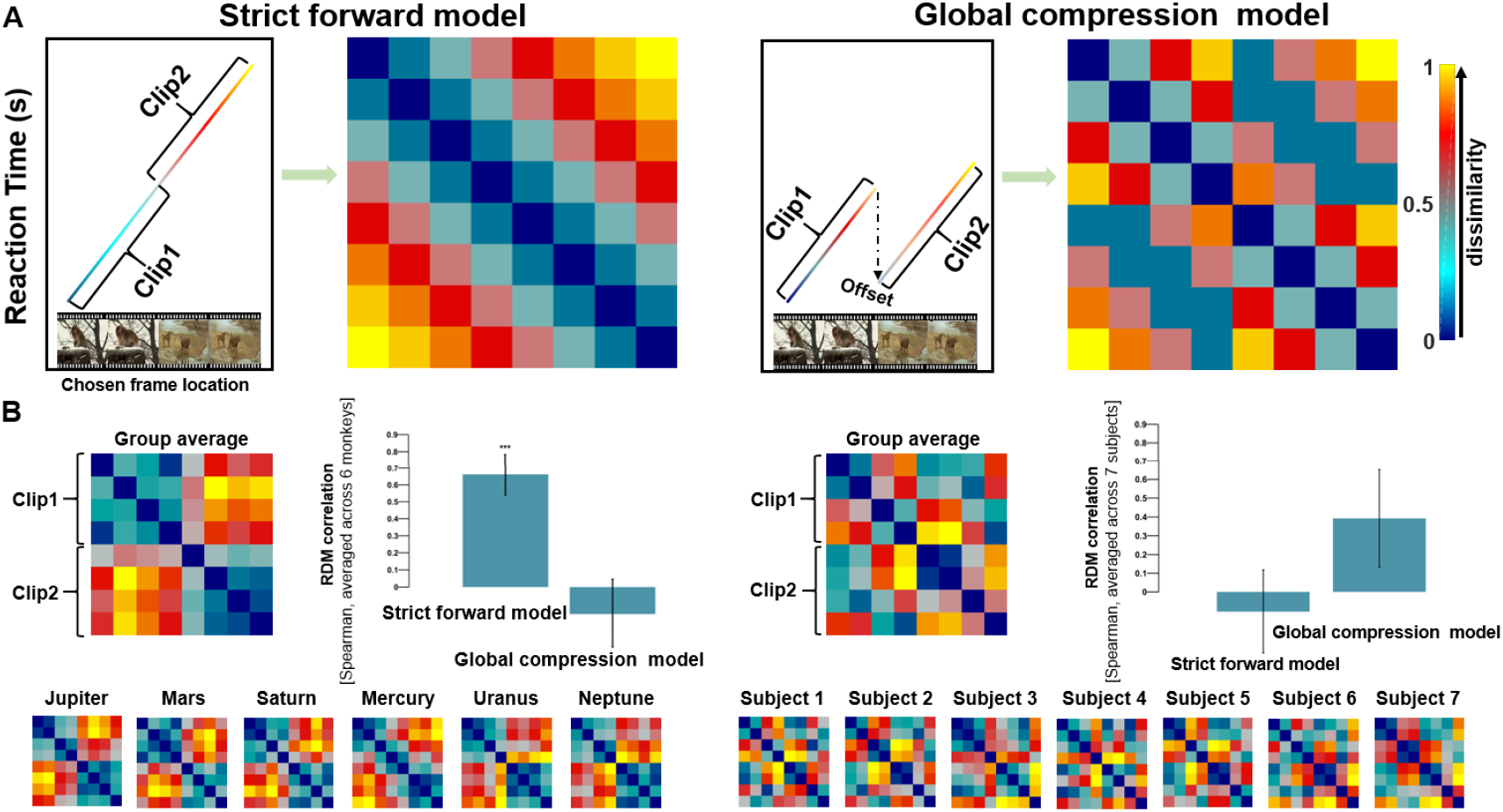
Model comparison using representative similarity analysis. **(A)** Visualization of two candidate models as RDMs. Pattern of reaction time (rank-transformed values) as a function of increasing chosen frame location underlying the two hypothetical models. The colors of Clip 1 and Clip 2 evolve increasingly with the temporal progression of the video (left); and their respective hypothetical RDMs (right). The reduction in RT indicated by an arrow between Clip 1 and Clip 2 is defined as “offset”; the magnitude of such “offset” is arbitrary but see further analysis in Figure 4. **(B)** We segmented the videos into eight equal segments and the RDMs contain pairwise Euclidean distances between these different segments for the group average (Monkeys: left; humans: right) and each individual separately (Monkeys: left bottom; humans: right bottom). RDM correlation tests between behavioral RDMs and two candidate RDMs show that the monkeys replay the footage using a Strict forward strategy and show little evidence for the Global compression strategy. Humans show an opposite pattern from the macaques. In the humans (right panels), the Global compression model shows a higher correlation with behavioral RDM (marginally insignificant *r* = 0.39, *P* = 0.069) than with the Strict forward model (*r* = −0.11, *P* = 0.703). Error bars indicate the standard error of the mean based on 100 iterations of randomization. *P* values are FDR-corrected (*** denotes *P* < 0.001).

### Factors modulating the model: “offsets” for search and memory-search RT slope

We defined the reduced RT to initiate replay for Clip 2 as “offsets” in initiating search in Clip 2 by skipping the non-informative Clip 1 (Figure 3A). With respect to the detailed differences between the two models, one may wonder whether and how the “offsets” between Clip 1 and Clip 2 might influence the results. Especially for the Global compression model, changes of this parameter will cause changes in the RDMs. To address this concern, we simulated an array of RDMs by systemically varying the offset parameter and produced 11 hypothetical models ranging from an absolute Global compression model (model 1, most left in Figure 4A), to a Strict-forward model (model 6, middle in Figure 4A), and beyond (7^th^ to 11^th^ models, right in Figure 4A). We then tested each individual monkeys’ data with each of these 11 models. The results show that the Spearman correlation values between the monkey’s data and hypothetical RDMs reach an asymptote of around *r* = 0.8 as the offset parameter tend to zero, and notably, the correlation values only improve minimally with increasing offsets (Figure 4B, with individual’s RT RDM displayed as insets). These suggest that the monkeys have processed the video as a holistic chunk of information rather than taking advantage of skipping the non-informative first clip when the two probe frames were in Clip 2. For comparison, we also tested human participant’s data against each of these 11 hypothetical models and found a completely opposite pattern in the humans (Figure 4C, with individual’s RT RDM displayed as insets). Taken together, we reveal a discrepancy between human and macaque performance in terms of their ability to compress past (irrelevant) information during TOJ.

**Figure 4.**
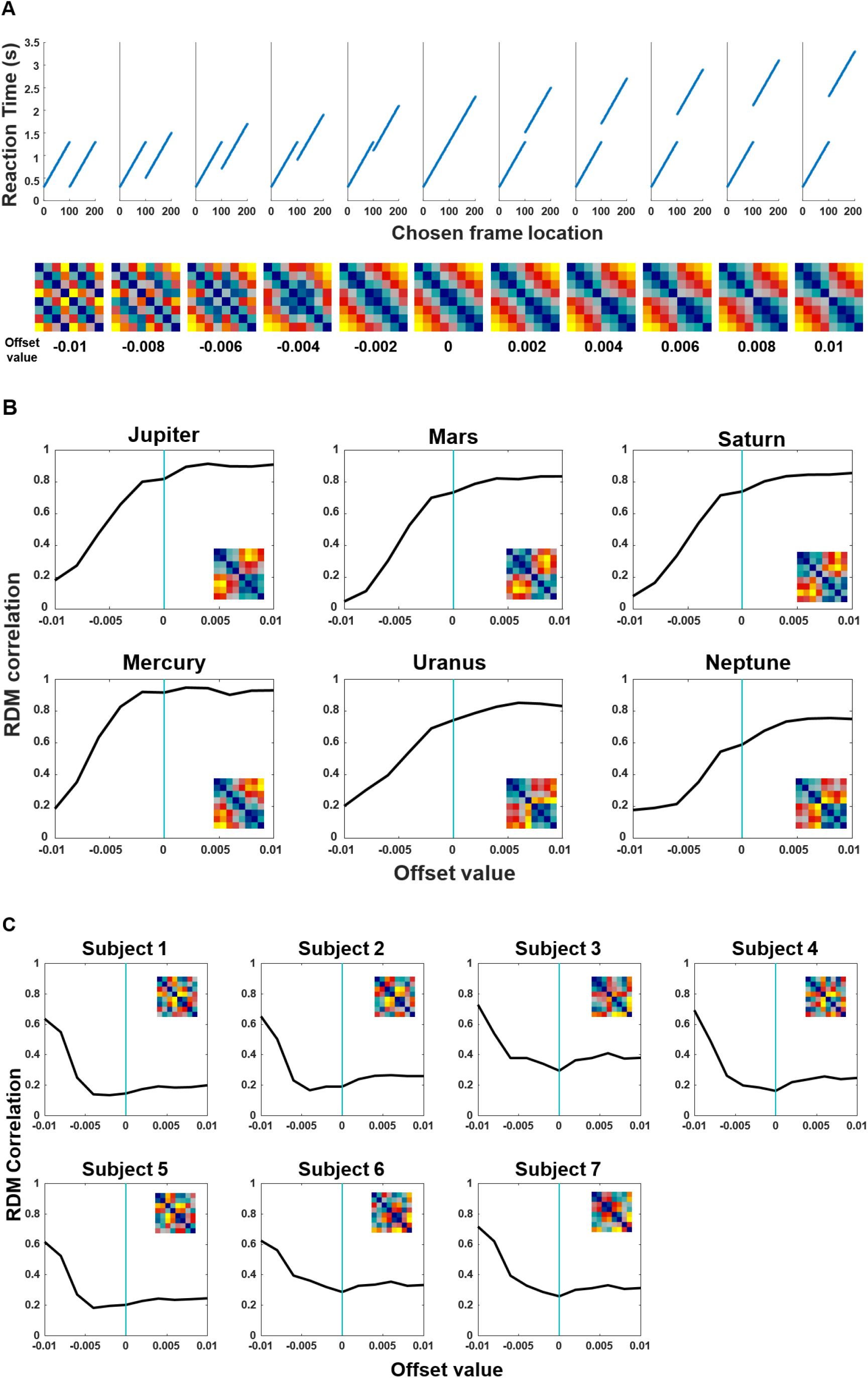
Strict forward model provides better fit to the RT data in monkeys but not in humans. **(A)** “Offsets” are defined as the magnitude of reduced RT when the frames were in Clip 2. 11 hypothetical models with their reaction time patterns (top) and RDMs (bottom). We systemically varied the “offset” parameter while keeping a constant slope. These models progressively range from an absolute Global compression model (model 1, most left), to a Strict-forward model (model 6, middle), and beyond (7^th^ to 11^th^ models, right). The numerals below the RDMs denote the magnitude of the respective offsets. **(B)** Each monkey’s data is tested against each of these 11 hypothetical models. The Spearman correlations increase as a function of offset magnitude between Clip 1 and Clip 2 until reaching an asymptote when the offset value is around zero, which corresponds to the Strict forward model (model 6 in Figure 4A; see also Figure 3). Individuals’ RT RDMs are shown in insets. **(C)** Each human participant’s data is also tested against each of these 11 hypothetical models. The Spearman correlations decrease as a function of offset magnitude between Clip 1 and Clip 2 until reaching an asymptote when the offset value is around zero. This confirms the hypothetical discrepancy between the two species (see also Figure 3B).

We have considered a further factor – slope of RT/chosen frame location – in our model, which might be relevant for the question at hand. To take negative slopes into consideration, we calculated RDM based on the relative differences of each pair of segments. Then, we transformed the vectors “offset” and “slope of RT/chosen frame location” into two arrays and displayed them as three-dimensional mesh/surface plots. In Figure 5, in addition to “offsets”, we showed that the correlation between model and monkeys’ data also increase as the slope of RT/chosen frame location increases. In contrast, the correlation values for human data seem to be driven by the “offsets” (cf. Figure 4C), and in terms of “slope” going in an opposite direction from the monkeys (cf. Figure 2A-B). This visualization converges with other current findings that the slope of RT/chosen frame location does also matter to highlight the cross-species differences (e.g., by contrasting Figure 1C vs. 1D).

**Figure 5.**
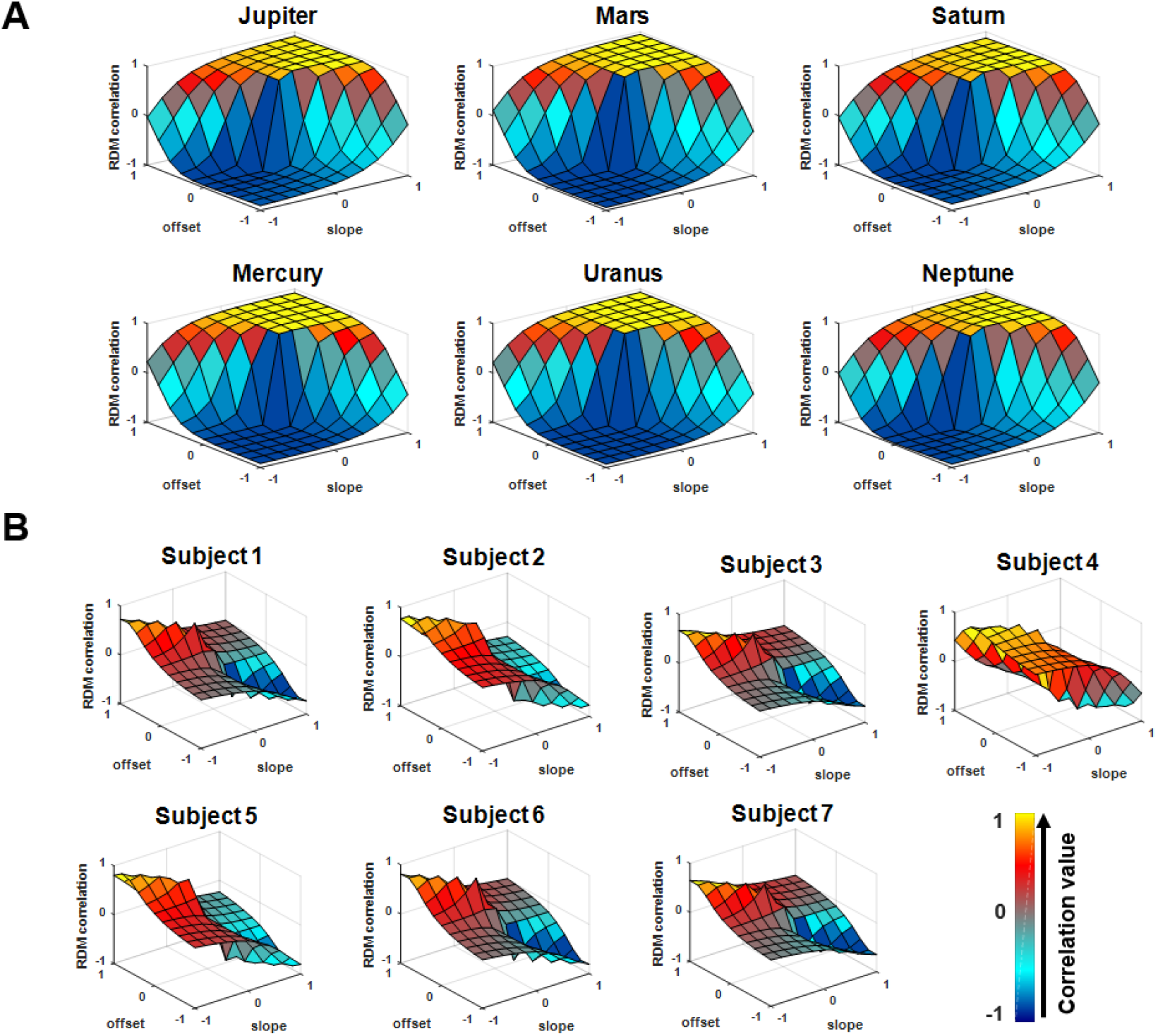
Visualization for RDM correlation as modulated by “offset” and “slope” in three-dimensional space (slope, offset, RDM correlation). **(A)** Monkeys data **(B)** Human participants data.

### Context changes (event boundary) increase rate of rise in decision information

We have thus far focused on how the monkeys retrieve the order of frames when information was equated within contexts, but how contextual changes would aid temporal order judgement processes remains to be examined. It was evident that the monkeys retrieved the temporal order of frames with numerally different speeds for the three trial-types, across- vs. within-Clip1 vs. within-Clip2: *F* (2, 15) = 2.32, *P* = 0.132 (Figure 1B, right). Thus, we then compared the latency distribution of within-context and across-context conditions and hypothesized that a context shift would change the rate of rise of information accumulation (shift model) without altering the decision threshold (swivel model) with the Linear Approach to Threshold with Ergodic Rate (LATER) model (Figure 6A). We compared across-context and within-context trials specifically and fitted the two types of LATER models separately on each monkey’s data (Figure 6B), together with an unconstrained model, which supposes the reaction times of two conditions are independent from each other, and a null model, which assumes there is no effect of manipulation. Using the Bayesian information criterion (BIC) as an index of model comparison, the results consistently indicate that the shift model is better than the swivel model in all six monkeys (range of ΔBIC = [14.57, 300.07]; Table S1). These results further indicate that contextual changes do not alter the judgement threshold for decisions (no evidence for a swivel pattern). Within a drift diffusion model framework, the results suggest that monkeys accumulate information for memory decisions at a faster rate when the frames were extracted from two different clips than for frames that were extracted from the same-context clip, signifying a boundary effect in memory in macaque monkeys which parallel closely those established findings in the humans (*17*). These results suggest that the monkeys adopt a forward search for targets among linearly ordered memory traces and they reach their memory decision threshold more quickly when probe frames are extracted from the two different contexts (across-context condition).

**Figure 6.**
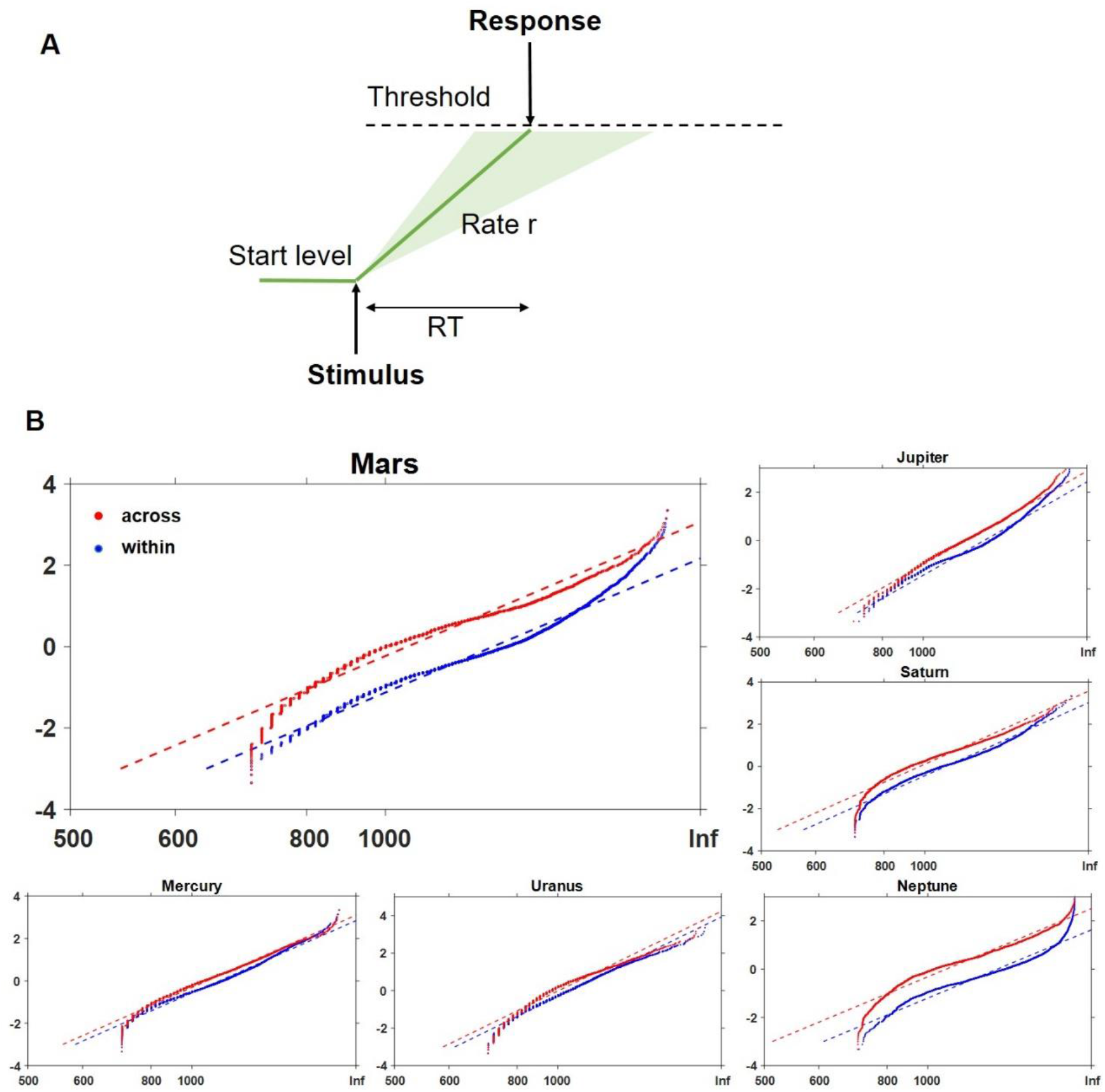
LATER model fitting of RT in across-context and within-context condition. **(A)** The LATER model cartoon depicts that a decision signal triggered by a stimulus rises from its start level, at a rate of information accumulation *r*, to the threshold. Once it reaches the threshold, a decision is initiated. The rate of rise *r* varies from trial to trial, obeying the Gaussian distribution (variation denoted as green shaded area). **(B)** Contextual change effect on the distribution of response latency for the monkeys; data from Monkey “Mars” was chosen for larger display. The red and blue dash lines show the best fits (maximum likelihood) of across-context trials and within-context trials respectively.

### Confirmatory GLMs for the putative patterns

In order to verify whether the effects are not attributed to the basic stimulus features such as perceptual differences inherent in the across-context condition. We then performed several generalized linear models to quantify the effect sizes of several principal variables. In within-context condition, given that the monkeys would replay their experience to judge the relative temporal order of probe frames (“replay hypothesis”), we used temporal characteristics of probe frames, as represented by chosen frame location (or temporal similarity, which is essentially an inverse of frame location) as the independent variables. In across-context condition, we included a perceptual similarity measure based on feature points extracted by the SURF algorithm (SURF similarity, Figure S2) into the generalized linear model to reflect the extent to which the monkeys could capitalize on using contextual boundaries for TOJ judgment. The within-context GLM shows that monkey’s RT was indeed significantly faster when the probe frame was located earlier in the video, *P* < 0.001 (or in equivalent terms, when two frames were temporally closer, *P* = 0.004), confirming our main finding that the monkeys adopt a forward scanning strategy for information retrieval (Figure 7 left). In contrast, the across-context GLM shows that there was not any significant effect of the chosen frame location on RT. Rather, the monkeys retrieve their memories significantly faster for probe frames that are contextually (or perceptually) distinct, *P* = 0.004 (Figure 7 right). These results are fully replicated when several more regressors which can potentially affect RT are additionally entered into the design matrix (Figure S3).

**Figure 7.**
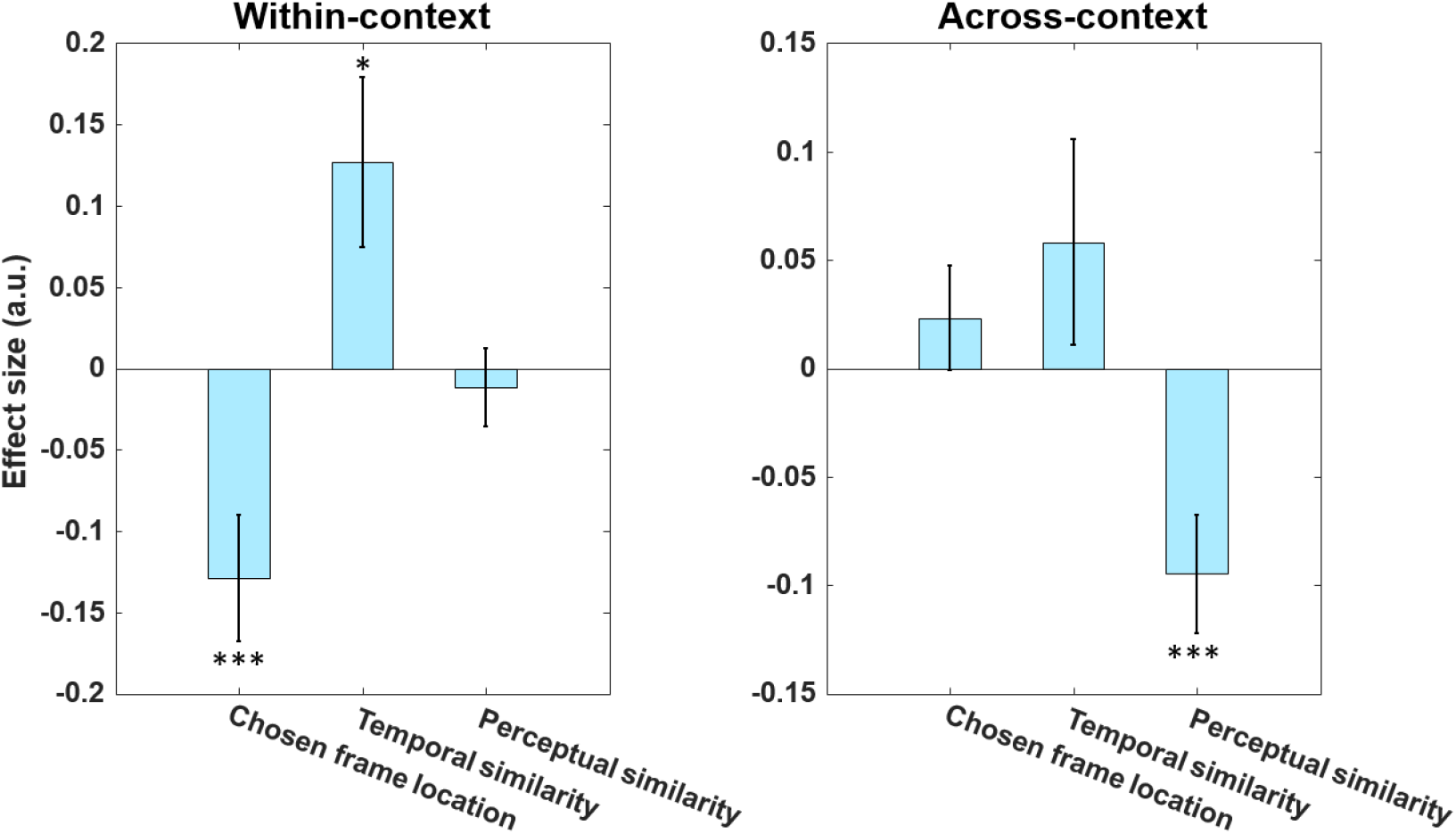
GLM results on the effects of model variables on reciprocal latency for within-context and across-context condition. The chosen frame location and the temporal similarity are variables associated with the replay hypothesis, whereas the perceptual similarity is with the context hypothesis. In within-context condition, the chosen frame location and the temporal similarity both show significant effects on RT, *P* < 0.001 and *P* = 0.004 respectively, wheras the perceptual similarity does not, *P* = 0.844. We found an opposite effect pattern in across-context condition. The perceptual similarity shows a significant effect on RT, (*P* = 0.004), while the chosen frame location (*P* = 0.796) and the temporal similarity (*P* = 0.446) do not. Effect size (in arbitray unit, a.u.) denote the beta values. Error bars indicate SEM. *** denotes *P* < 0.001, ** denotes *P* < 0.01, * denotes *P* < 0.05. See also Figure S3 for a GLM analysis including a full range of variables.

## DISCUSSION

In light of recent reports on neural correlates underlying how humans and rodents reply their past experiences (*11*, *12*, *14*, *18*), here, we demonstrate parallel behavioral findings in macaque monkeys with dynamic cinematic material. Previous reports of macaques succeeding in temporal order judgement indicated their ability in remembering the order of events (*1*, *5*, *19*) and even meta-cognitively monitor the quality of representations of temporal relations among item images (*20*). For example, Orlov et al. (*8*) suggested that monkeys can categorize stimuli by their ordinal number to aid recall of order, and Templer et al. showed that monkeys retrieve the temporal order information based on the order of events rather than elapsed time (*1*). One possible common mechanism underlying these performances is that monkeys use a forward search to identify targets in memory representation (*2*). Taking advantage of the latency data obtained during TOJ on naturalistic materials, we provide new behavioral evidence in support of the hypothesis proposed by Gower (*2*) that the monkeys can replay their memory in a serial forward manner. Our analysis further clarifies that this replay process is conducted in a time-compressed manner. Notwithstanding task differences, both humans and macaques execute retrieval with forward replay with a comparable compression factor (factors of ~11 in macaques vs. ~13 in humans, cf. (*12*), but see also (*13*)).

Despite the cross-species similarity, our revelation of a critical discrepancy between humans and macaques carries an important theoretical implication: humans can do both local and global compression whereas monkeys are not able to attain global compression. The implication is that mental time travel is not all-or-none. There could be multiple layers underpinning the concept of mental time travel, which entail the ability to relive the past (*21*, *22*) and skipping over unimportant information (*12*). The latter aspect allows humans to recall our memories flexibly far into our past and suggest powerful computational efficiencies that may facilitate memory storage and recall. In contrast, we did not find any evidence in the macaques for their ability of making use of salient boundary cues to skip unimportant details. It thus remains unknown in the current experiment how far back in time monkeys can replay their memories. It has been recently shown that humans can spontaneously replay experience based on learned structures, with fast structural generalization to new experiences by representing structural information in a format that is independent from its sensory consequences (*11*). The lack of global compression in the monkeys of their video experience implies that the monkeys might not be able to use factorized representations to allow components to be recombined in more ways than were experienced (*23*).

A further caveat is that our monkeys perform their replay-like recall of the videos as an effortful operation to solve a TOJ task whereas most of the extant studies on replay focus on spontaneous replay patterns at rest (*11*), during sleep (*24*, *25*) or during task-free state (*26*, *27*). While we have argued for the presence of replay-like patterns in the monkey during TOJ, we are aware that replay is a neural phenomenon supported by activity of individual neurons and implicates offline reactivation of sequences of hippocampal place cells that reflect past and future trajectories (*28*–*30*). However, by establishing that neither the number of intervening frames nor passage of time per se determines the RT pattern (probably a mixed effect resultant from a combination of both), we ruled out order- or positional-memory as the underlying mechanism supporting TOJ in this task. Our results combined thus provide a novel connection between various kinds of replay(-like) behaviors linking between rodents and humans and provide a primate model for neuronal investigation.

We observed one further interesting feature here that the monkeys are able to detect contextual changes to facilitate TOJ. We show that the expedited TOJ in across-context condition was facilitated by contextual details, which in turn results in an increased rate of rise of signal towards memory decision in a drift-diffusion process. Humans studies show that contextual changes lead to segmentation of ongoing information (*31*, *32*). Our results provide evidence consistent with event segmentation in the macaque monkeys and imply that macaque monkeys might be capable of parsing the footage by contextual information, akin to what has been shown in humans (*33*–*36*) and rodents (*37*).

Memory replay is an elaborate mental process and our demonstration of a time-compressed, forward replay-like pattern in the macaque monkeys provides insights into mapping the mechanisms and evolution of episodic memory in our lineage. There are however limitations in their ability in compressing the experienced past globally, which somehow indicates a form of primordial mnemonic rigidity. This cognitive discrepancy in our lineage should be further elucidated via electrophysiological or neuroimaging methods probing into the MTL (*18*, *26*, *38*, *39*) and the neocortices (*16*).

## METHODS

### Task performance

The six monkeys performed the task with a significantly above chance level with an overall accuracy at 67.9 ± 1.5% (mean ± SD) and with an above chance accuracy for within-context trials, *t*5=14.35, *p* < 0.001. The human participants performed the task on average at 92.7 ± 1.2%.

### Subjects

#### Macaque monkeys

Six male rhesus macaques (*Macaca mulatta*) (5.49 ±0.5kg) with a mean age of 3.5 years at the start of testing participated in this study. They were initially housed in a group of 6 in a specially built spacious enclosure (max capacity = 12-16 adults) with enrichment elements such as a swing and climbing structures until the present study began. The monkeys were then housed in pairs during experimentation period according to their social hierarchy and temperament. They are fed twice a day with portions of 180-g monkey chow and pieces of fruits (8:30am/4:00pm). Water is available *ad libitum* except on experimental days. They are routinely offered treats such as peanuts, raisins and various kinds of seeds in their home cage for forage purpose. The monkeys were procured from a nationally accredited colony located in Beijing outskirts, where the monkeys were bred and reared. The animals are thus ecologically naive to the natural wilderness and should not have had any previous encounter with other creatures except humans and their companion. The room wherein they are housed is operated with an automated 12:12 (7am/7pm) light-dark cycle and kept within temperate around 18-23°C and humidity of 60-80%.

#### Human subjects

Seven participants (mean age = 19.57 ± 1.13, 6 female) took part in the experiment. The participants were recruited from the undergraduate population in East China Normal University. The participants provided informed consent and were compensated 400 RMB for their time. We used the 6 unique video-trials sets and TOJ frames (one unique set per monkey) correspondingly for the human subjects (subject 7 re-used set 1).

The experimental protocol was approved by the Institutional Animal Care and Use Committee (permission code: M020150902 & M020150902-2018) and the University Committee on Human Research Protection (permission code: HR 023-2017) at East China Normal University. All experimental protocols and animal welfare adhered with the “NIH Guidelines for the Care and Use of Laboratory Animals”.

### Apparatus and testing cubicle

The testing was conducted in an automated test apparatus controlled by two Windows computers (OptiPlex 3020, Dell). The subject sat, head-unrestrained, in a wheeled, specially-made Plexiglas monkey chair (29.4cm × 30.8cm × 55cm) fixed in position in front of a 17-inch infrared touch-sensitive screen (An-210W02CM, Shenzhen Anmite Technology Co., Ltd., China) with a refresh rate at 60 Hz. The distance between the subjects’ head and the screen was kept at ~20 cm. The touch-sensitive screen was mounted firmly on a custom-made metal frame (18.5cm × 53.2cm) on a large platform (100cm × 150cm × 76cm). Water reward delivery was controlled by an automated water-delivery rewarding system (5-RLD-D1, Crist Instrument Co., Inc., U.S.) and each delivery was accompanied by an audible click. An infrared camera and video recording system (EZVIZ-C2C, Hangzhou Ezviz Network Co., Ltd., China) allowed the subject to be monitored while it was engaged in the task. The entire apparatus was housed in a sound-proof experimental cubicle that was dark apart from the background illumination from the touch screen.

### Source of video materials and preparation

A collection of documentary films on wild creature was gathered from YouTube. The films were *Monkey Kingdom* (Disney), *Monkey Planet* (Episode 1 – 3; BBC), *Planet Earth* (Episode 1 – 11; BBC), *Life* (Episode 1 – 10; BBC), and *Snow Monkey* (PBS Nature). In total 28 hours of footage was gathered. We applied Video Studio X8 (Core Corporation) to parse the footage by camera-cuts into smaller segments. Experimenters then applied the following criteria to manually edit out ~ 2500 unique clips: 1) the clip must contain a continuous flow of depiction of events (i.e., no scene transition); 2) at least one living creature must be included; 3a) at least one of the animals contained must be in obvious motion; 3b) the trajectories of these motion must be unidirectional (i.e., no back and forth motion of the same subject); 4) clips with snakes were discarded. From this library, we then selected 2000 clips (all of 4 s – 6 s) for the final test and a small number of additional clips were also prepared for the training stages. For each individual monkey, the videos assigned to be in an experimental condition would not be used/shown in another condition so that a particular monkey will only view that video repeatedly under the same condition across exposure.

### Task and experimental procedure

We combined naturalistic material with a temporal order judgement paradigm that is widely used in episodic memory research (*1*, *40*, *41*). In each trial, the monkey initiated a trial by pressing a colored rectangle in the center of the screen (0.15 ml water). An 8-12 s video (consisting of two 4-6 s clips) was then presented (0.15 ml water), and following a 2-s retention delay, two frames extracted from the video were displayed bilaterally on the screen for TOJ. The monkeys were trained to choose the frame that was shown earlier in the video (see **Movie S1**). A touch to the target frame resulted in 1.5 ml water as reward, removed the foil frame, and the target frame would remain alone for 5 s as positive feedback. A touch to the foil frame removed both frames from the screen and blanked the screen for 20 s without water delivery. Since the monkeys could self-start the trials, we did not set an explicit inter-trial interval. Correction trial procedures were not used in the main test.

We collected 50 daily sessions of data. Each session contained 100 trials, giving us a 5000 trials per monkey. The 5000 trials contained a break-down of four factors: Boundary (Within vs. Across), Play Order (Normal vs. Reverse), Temporal distance (TDs, 25 levels), and Exposure (R1 – R5), giving out a 2 * 2 * 25 * 5 within-subject design. The two frames for TOJ could be extracted from same clip (Within) or distinct clips (Across). The 25 levels of TDs ranged between a minimum of 1000 ms (equivalent of 25 frames) and a maximum of 3880 ms (equivalent of 97 frames). Each TD level increased progressively with 3 frames each step. The 10 lists contained 5 lists of primate animals and 5 lists of non-primate animals (Category: Primate/Non-Primate). The 6 monkeys were counter-balanced in their order of receiving the two kinds of material: three monkeys (Jupiter, Mars, and Saturn) were tested first on Non-Primate lists, whereas the other three (Mercury, Neptune, and Uranus) were tested first on Primate lists. The whole experiment lasted for 68 days. There were three blocks of reset (3 days, 9 days and 6 days) in-between testing days. The task was programmed in in PsychoPy2 implemented in Python.

### Data analysis, temporal and perceptual similarity, and model comparison

#### Representational similarity analysis (RSA)

To create RDMs for the representational pattern of response time, we evenly divided each video into eight segments based on chosen frame location and averaged the response times within each segment for each monkey individually. For each matrix, we then computed the Euclidean distance of average response time for each pair of the segments. In order to show the relative distance among all pairs of segments intuitionally, we further transformed the matrix by replacing each element with the rank number in the distribution of all the elements, and linearly scaled into [0,1]. To compare similarity between response times patterns and the candidate models, we computed Spearman correlation between the model and group averaged RDMs with 100 randomized iterations. The statistical threshold was set at *p* < 0.05 (FDR corrected).

#### LATER (linear approach to threshold with ergodic rate) modelling

The observations that the brain needs longer time than it requires for nerves to transport information and that trial-by-trial RTs vary considerably have stimulated researchers to make use of distribution of reaction times for examining mental processes (*37*). Latency, as an indicator of decision processes, provides a source of insight into the underlying decision mechanisms (*34*–*36*). The Linear Approach to Threshold with Ergodic Rate (LATER) model is a widely used model that taps into these processes (*42*). The LATER model stipulates that the winner signals that reach threshold faster would trigger the decision, resulting in shorter latency. Accordingly, changing the rate of rise would cause the line to “shift” along the abscissa without changing its slope. In contrast, lines plotted by latency distribution would “swivel” around an intercept when the threshold changes (*43*) (Figure 4A). To explore the underlying mechanisms of different experiment conditions, we plotted the reciprocal of reaction time as a function of their z-scores, on account for making the distributions follow Gaussian distribution (*44*). Hence, we fitted the main component for each condition. The main component is a ramp to the threshold with rate of rise *r*. The distance between start level S_0_ and threshold S_T_ is defined as *θ*, and rate of rise *r* follows a Gaussian distribution of mean, *μ*, and standard deviation, *σ1*.

For model comparison, we fitted four different models to the data. The “null” model fits reaction times of both conditions with the same parameters, implying no effect of manipulation. The “Unconstrained” model set all the parameters to be free, which supposes the reaction times of two conditions are independent from each other. The “Shift” model only allows the slope of main component *μ* to change according different conditions, on the assumption that the manipulation will change the rate of rise. The “Swivel” model only allows *θ* to change according different conditions, on the assumption that the subject set different thresholds for different conditions.

#### Generalized linear models (GLM)

We ran generalized linear models to compare the effect sizes of independent variables to the dependent variable. The mean (*μ*) of the outcome distribution Y depends on the independent variables X, through the following formula:

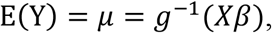

where Y is a distribution of outcomes, β is unknown parameters to be estimated, *g* is a link function (Gaussian function). The dependent variable (Y) is reciprocal latency, and independent variables are as follows: a binary regressor indicating whether the video includes primate content or not, a binary regressor indicating that a video is played forward or backward, the five repetitions of the video-trials, physical location of the selected probe (left or right), time elapsed within session, chosen frame location, temporal similarity, perceptual similarity (SURF), and temporal distance. To reduce the effect of behavioral noise of monkeys on the analysis, we included only correct trials in these GLM analyses and excluded trials with RT longer than 10 s or shorter than 0.7 s.

#### Model comparison

To obtain the best fit among these models, we used Bayesian Information Criterion (BIC) as a criterion for model selection among these four models. The formula for BIC is: BIC = −2(logL) + numParam* log(numObs), where L is the maximum likelihood for the model, numParam and numObs represent the number of free parameters and the number of samples respectively. We computed ΔBIC as the strength of the evidence, which indicates to what extent the selected model is superior to other models. Different ranges of ΔBIC show different level of evidence: The value of ΔBIC larger than 2 shows a positive evidence, the value of ΔBIC larger than 6 indicates a strong evidence (*56*).

#### Temporal similarity

For each trial, we calculated temporal similarity (TS) as an index of the discriminability of probe frames. Temporal similarity between two probe frames extracted from the video is calculated by the ratio of two frames’ temporal separation between their occurrence in the video and the time of testing. Temporal similarity of any two memory traces can be calculated as: TS = delay2 / delay1, where delay2 < delay1 (*45*).

#### Perceptual similarity

RGB-histogram is computed as the Sum-of Square-Difference (SSD) error between image pairs for the three color channels (RGB). For each color channel the intensity values range from 0 to 255 (i.e., 256 bins), we first computed the total number of pixels at each intensity value and then computed the SSD for all 256 bins for each image pair. The smaller the value of the SSD, the more similar 658 the two images (image pair) was. In the case of the HOG similarity, we constructed a histogram of directions of gradient over fixed sized grid across the entire image. A vector is generated from each grid cell and correlated with HOG features from another image. Besides, Speeded Up Robust Features (SURF) (*46*) uses Box Filter which is calculated in parallel using integral images (*47*) to approximate Laplacian-of-Gaussian (LoG). Wavelet responses in both horizontal and vertical directions are used to assign orientation in SURF. The first step in SURF consists of fixing a reproducible orientation based on information from a circular region around the interest point. A descriptor vector is generated around the interest point using integral image, which is compared to descriptor vectors extracted from a compared image to find a match. The Euclidean distance is used to measure the similarity between two descriptor vectors from two images.

#### Data preprocessing

Trials with RT longer than 10 s (1.45%) or faster than 0.7 s (2.47%) were excluded from the analyses. Both correct and incorrect trials were entered for all analyses, except those reported in the GLMs. For Neptune, data of one day of primate list 5 (exposure 4) and two days of primate lists 4 and 5 (exposure 5) were lost due to machine breakdown and thus only 4,700 trials are included for Neptune.

## Author contributions

Conceptualization, S.Z., L.W. and S.C.K.; Methodology, S.Z., L.W., J.S., S.W.L., K.A., and S.C.K.; Investigation, S.Z., L.W., and Y.C.; Data interpretation: S.Z., L.W., J.S., S.W.L., Y-d.Z., and S.C.K.; Writing – Original Draft, S.Z. and S.C.K.; Writing – Review & Editing, S.Z. and S.C.K.; Funding Acquisition, Y-d.Z., and S.C.K.; Supervision, S.C.K.

## Data and materials availability

All data is available at Dryad (doi:10.5061/dryad.3r2280gcc) and codes are available upon request.

## Supplemental Information

**Figure S1.**
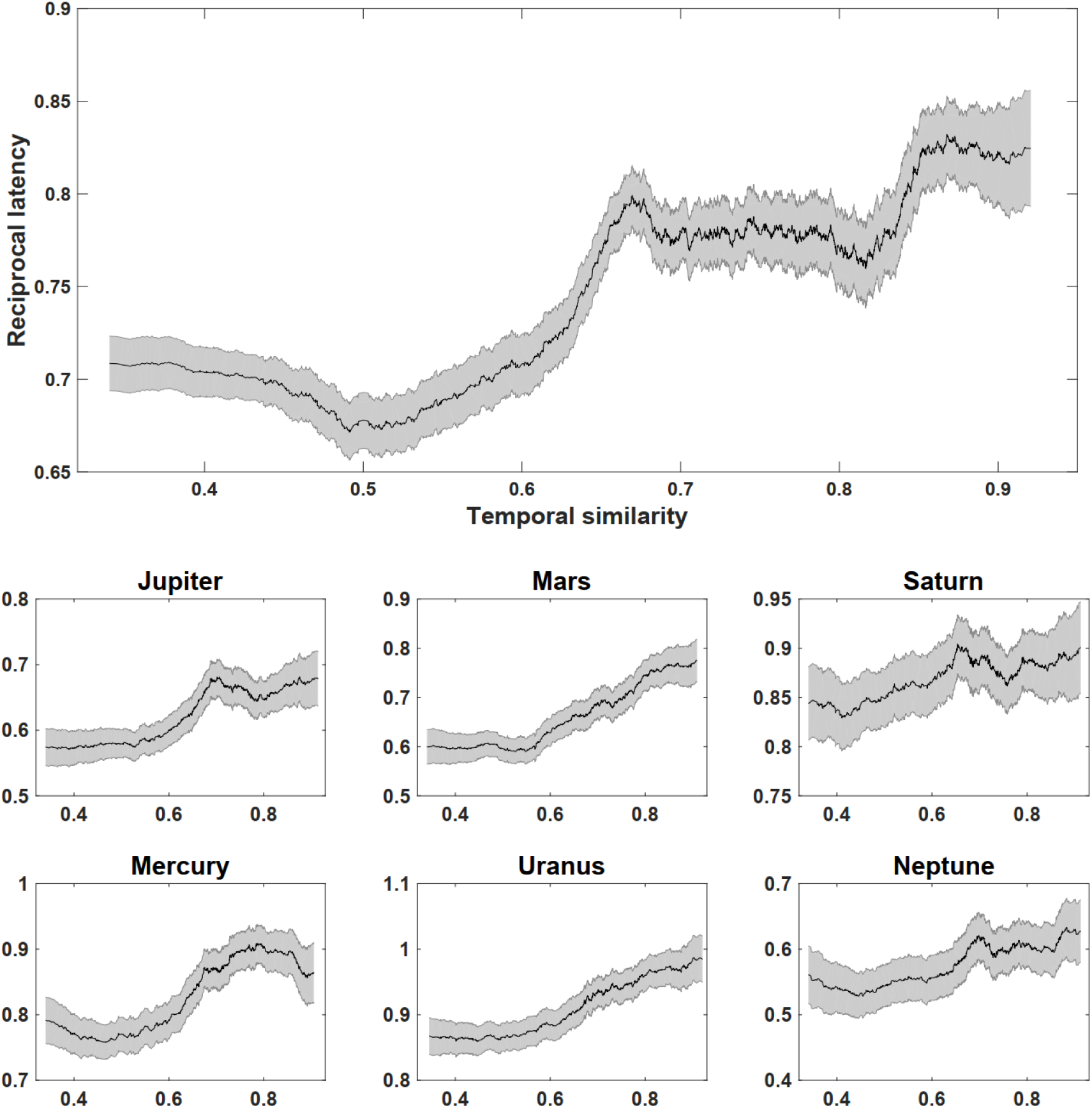
Relationship between temporal similarity and reciprocal latency for within-context trials. Reciprocal latency as a function of temporal similarity for the average of all monkeys (upper panel) and individual monkeys (bottom panel). Temporal similarity between two frames is mathematically the inverse of their respective frame locations in the video, see Methods. **Related to** Figure 2A.

**Figure S2.**
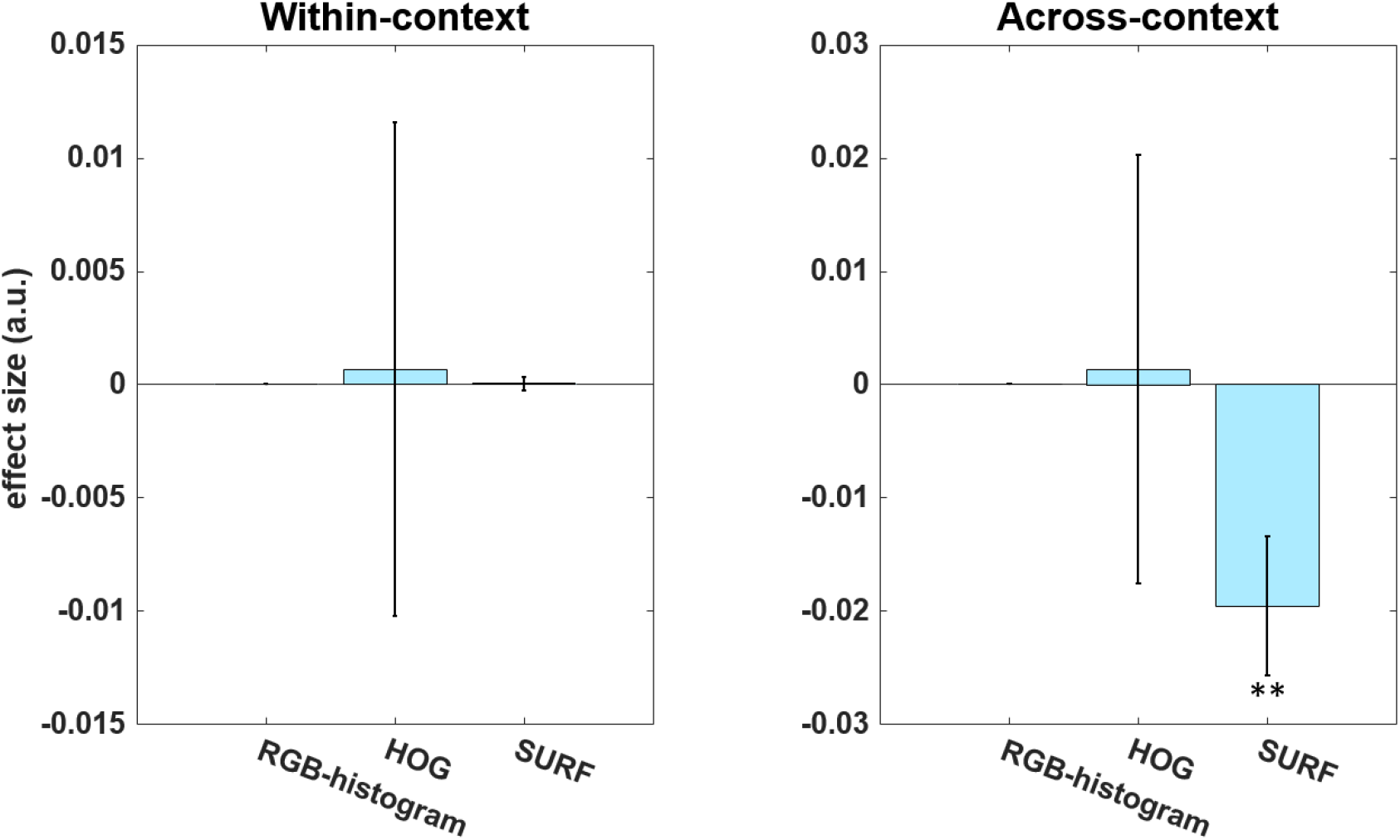
GLM results on the effects of image similarity measures on reciprocal latency for within-context and across-context condition. Difference of distribution in RGB- histogram, HOG similarity and SURF similarity were tested. The SURF similarity measure was significantly correlated with RT in the across-context condition (*P* = 0.0015). **Related to** Figure 7.

**Figure S3.**
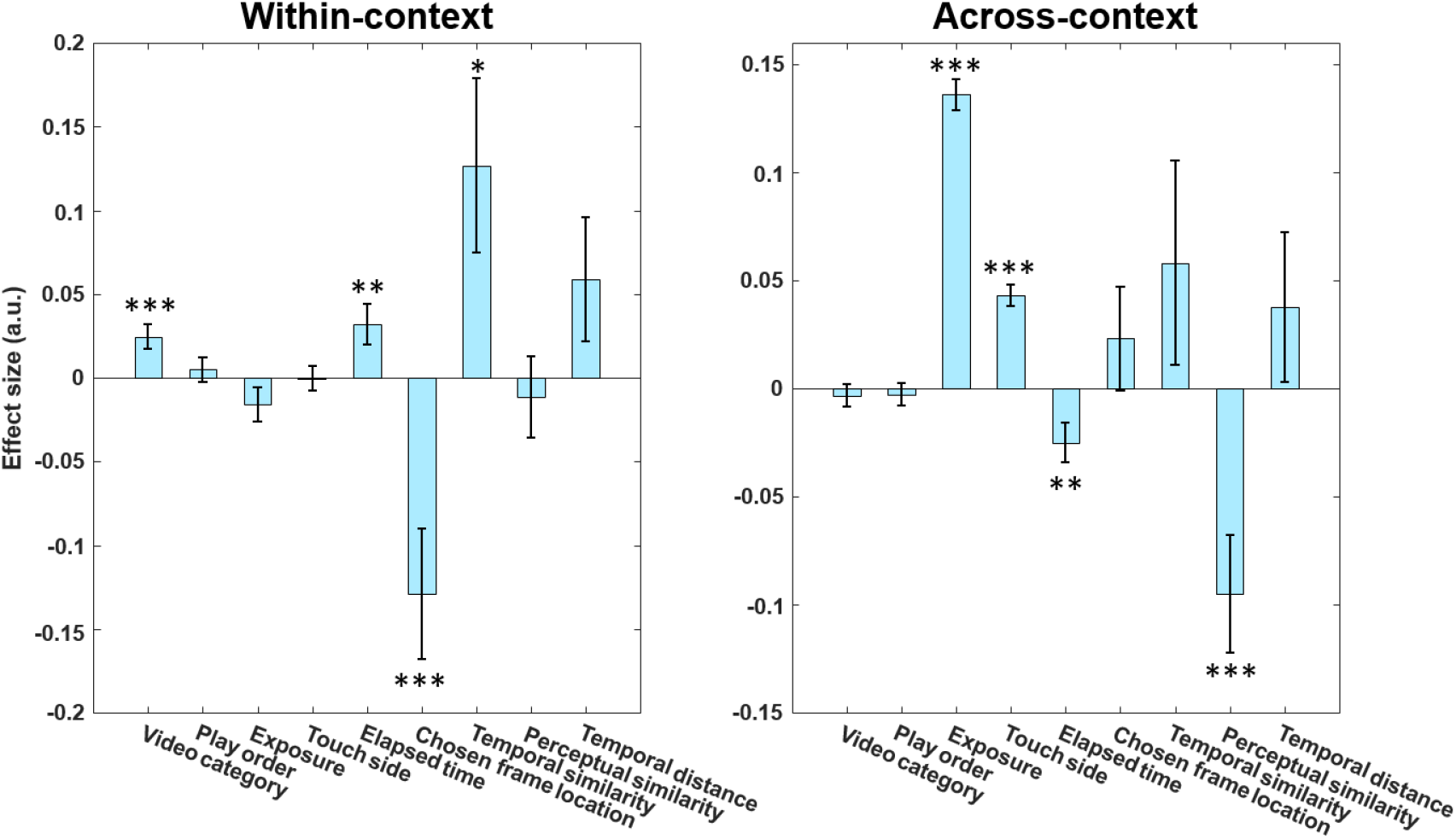
Full GLM analysis including a number of variables that might affect reciprocal latency. The nine regressors include a binary regressor indicating whether the video category is primate or non-primate (video category), a binary regressor indicating that a video is played forward or backward (play order), the repeated exposure of the trial (1 – 5) (exposure), the physical location of the selected probe on screen (left or right) (touch side), time elapsed within session (elapsed time; to rule out fatigue or attentional confounds), chosen frame location, temporal similarity, SURF similarity as a perceptual similarity measure (perceptual similarity) and temporal distance between two probe frames. The results confirm that chosen frame location is the most significant regressor in within-context trials while perceptual similarity is the most significant regressor in across-context trials. *** denotes *P* < 0.001, ** denotes *P* < 0.01, * denotes *P* < 0.05. **Related to** Figure 7.

**Table S1.** LATER model fitting results of six monkeys. For ease of comparison, we computed the respective ΔBIC to index the strength of evidence for each model. Note that the model with the lowest BIC is the winning model. In all 6 monkeys, the shift model was superior to the other three models. **Related to** Figure 6.

| Condition |  | BIC |  |  |  | delta_BIC |  |  |  |
| --- | --- | --- | --- | --- | --- | --- | --- | --- | --- |
|  |  | null | two fits | swivel | shift | null | two fits | swivel | shift |
| Across VS Within | Jupiter | -67157.39 | -67603.99 | -67543.25 | -67615.23 | 457.83 | 11.24 | 71.98 | 0 |
|  | Mars | -63948.06 | -65937.89 | -65769.84 | -65948.72 | 2000.65 | 10.82 | 178.88 | 0 |
|  | Saturn | -61307.14 | -61929.72 | -61853.77 | -61938.70 | 631.56 | 8.98 | 84.93 | 0 |
|  | Mercury | -63904.96 | -64047.31 | -64042.94 | -64057.52 | 152.56 | 10.20 | 14.57 | 0 |
|  | Uranus | -67652.66 | -67850.84 | -67841.34 | -67864.67 | 212.01 | 13.83 | 23.34 | 0 |
|  | Neptune | -56695.53 | -58327.76 | -58037.15 | -58337.23 | 1641.70 | 9.47 | 300.07 | 0 |

**Table S2.** For monkeys: One sample *t*-tests results of the slopes of RT as a function of temporal similarity. The three panels correspond to analyses performed using all trials (top), only correct trials (middle), and only incorrect trials (bottom). Same slope patterns were observed irrespective of correctness, in consistency with the analyses on slopes of RT as a function of chosen frame location for each monkey. **Related to** Table 1.

| Monkeys | Mean (SEM) | t-statistics | d.f. | p-value | Cohen's d | 95% confidence interval |  |
| --- | --- | --- | --- | --- | --- | --- | --- |
|  |  |  |  |  |  | Lower | Upper |
| Slope of reaction times/temporal similarity tested against zero (all trials) |  |  |  |  |  |  |  |
| Jupiter | -0.59 (1.04) | -3.99 | 49 | <0.001 | -0.56 | -0.88 | -0.29 |
| Mars | -1.15 (1.18) | -6.86 | 49 | <0.001 | -0.97 | -1.48 | -0.81 |
| Saturn | -0.24 (0.69) | -2.46 | 49 | 0.020 | -0.35 | -0.43 | -0.04 |
| Mercury | -0.52 (0.73) | -5.02 | 49 | <0.001 | -0.71 | -0.73 | -0.31 |
| Uranus | -0.44 (0.64) | -4.93 | 49 | <0.001 | -0.70 | -0.62 | -0.26 |
| Neptune | -1.03 (1.94) | -3.65 | 46 | <0.001 | -0.53 | -1.60 | -0.46 |
| Slope of reaction times/temporal similarity tested against zero (correct trials) |  |  |  |  |  |  |  |
| Jupiter | -0.63 (1.45) | -3.06 | 49 | 0.004 | -0.43 | -1.04 | -0.22 |
| Mars | -1.01 (1.44) | -4.98 | 49 | <0.001 | -0.70 | -1.42 | -0.60 |
| Saturn | -0.29 (0.80) | -2.58 | 49 | 0.013 | -0.36 | -0.52 | -0.06 |
| Mercury | -0.46 (0.89) | -3.64 | 49 | <0.001 | -0.51 | -0.71 | -0.20 |
| Uranus | -0.46 (0.70) | -4.64 | 49 | <0.001 | -0.66 | -0.66 | -0.26 |
| Neptune | -1.01 (2.69) | -2.56 | 46 | 0.014 | -0.37 | -1.80 | -0.22 |
| Slope of reaction times/temporal similarity tested against zero (Incorrect trials) |  |  |  |  |  |  |  |
| Jupiter | -0.63 (1.40) | -3.17 | 49 | 0.003 | -0.45 | -1.02 | -0.23 |
| Mars | -1.31 (1.70) | -5.43 | 49 | <0.001 | -0.77 | -1.79 | -0.82 |
| Saturn | -0.25 (0.28) | -1.39 | 49 | 0.169 | -0.20 | -0.61 | 0.11 |
| Mercury | -0.55 (1.40) | -2.81 | 49 | 0.007 | -0.40 | -0.95 | -0.16 |
| Uranus | -0.42 (0.90) | -3.29 | 49 | 0.002 | -0.47 | -0.68 | -0.16 |
| Neptune | -1.27 (3.42) | -2.55 | 46 | 0.014 | -0.37 | -2.27 | -0.27 |

**Table S3.** For human participants: One sample *t*-tests results of the slopes of RT as a function of temporal similarity (upper panel) and the slopes of RT as a function of chosen frame location (bottom panel) against zero. **Related to** Figure 1D.

| Subjects | Mean (SEM) | t-statistics | d.f. | p-value | Cohen's d | 95% confidence interval |  |
| --- | --- | --- | --- | --- | --- | --- | --- |
|  |  |  |  |  |  | Lower | Upper |
| Slope of reaction times/temporal similarity tested against zero |  |  |  |  |  |  |  |
| Subject 1 | 1.54 (1.17) | 5.87 | 19 | <0.001 | 1.31 | 0.99 | 2.09 |
| Subject 2 | 2.10 (1.84) | 5.10 | 19 | <0.001 | 1.14 | 1.24 | 2.96 |
| Subject 3 | 2.28 (1.56) | 6.55 | 19 | <0.001 | 1.55 | 1.55 | 3.01 |
| Subject 4 | 2.12 (1.30) | 7.32 | 19 | <0.001 | 1.51 | 1.51 | 2.73 |
| Subject 5 | 1.01 (1.05) | 4.30 | 19 | <0.001 | 0.52 | 0.52 | 1.50 |
| Subject 6 | 0.51 (0.80) | 2.87 | 19 | 0.010 | 0.41 | 0.14 | 0.89 |
| Subject 7 | 1.31 (1.77) | 3.29 | 19 | 0.004 | 0.48 | 0.48 | 2.14 |
| Slope of reaction times/chosen frame location tested against zero |  |  |  |  |  |  |  |
| Subject 1 | 0.0004 (0.0038) | 0.44 | 19 | 0.667 | 0.10 | -0.0014 | 0.0021 |
| Subject 2 | -0.0005 (0.0034) | -0.71 | 19 | 0.487 | -0.16 | -0.0021 | 0.0010 |
| Subject 3 | -0.0027 (0.0040) | -2.99 | 19 | 0.008 | -0.67 | -0.0045 | -0.0008 |
| Subject 4 | -0.0018 (0.0024) | -3.35 | 19 | 0.003 | -0.75 | -0.0029 | -0.0007 |
| Subject 5 | -0.0001 (0.0027) | -0.09 | 19 | 0.931 | -0.02 | -0.0013 | 0.0012 |
| Subject 6 | -0.0003 (0.0020) | -0.65 | 19 | 0.521 | -0.15 | -0.0012 | 0.0006 |
| Subject 7 | -0.0009 (0.0047) | -0.84 | 19 | 0.410 | -0.19 | -0.0031 | 0.0013 |

